# Characterization of the Oral Bacteriome of the Healthy Lewis Rat

**DOI:** 10.1101/2023.06.22.546140

**Authors:** Sareda TJ Schramm, Emily DeCurtis, Sutton E. Wheelis, Ian Jorgeson, Danieli BC Rodrigues, Kelli Palmer

**Author notes:** co-corresponding authors Contact information: Danieli Rodrigues, Kelli Palmer.

## Abstract

Rodents are the most frequently utilized animals for *in vivo* research, and rats are the most abundant rodent used to study peri-implantitis disease and progression. However, there are very few studies available that elucidate the healthy oral microbiome of rats. The aim of this study was to characterize the healthy oral bacteriome of the male Lewis rat at two unique niches, the gums and first molar, by sequencing the 16S rDNA v1-v3 region. We collected the microbiota of 12 male Lewis rats at the toothless alveolar crest and the first molar, on both sides of the mouth. We identified an average of 20,820 sequences per sample. There were no significant differences between the rat groups or sample sites in the diversity (α or β) of the bacteriome. A total of 359 amplicon sequence variants (ASVs) were identified in the bacteriome, correlating to the phyla *Firmicutes*, *Actinobacteria*, *Proteobacteria*, and *Bacteroidetes*. Parallels exist at the genus level between human and rat oral bacteriomes, however, the overall diversity between comparable sites is lower in rats than humans. Similar findings were reached when using pooled or individual swab samples of rat mouths. This study characterizes two unique ecological niches within the healthy rat oral bacteriome and provides a baseline for comparison with future studies.

**IMPORTANCE:** Rats are frequently used for dental research, but little is known about their normal oral microbiota. Here, we use DNA sequence analysis to identify bacterial groups colonizing the rat mouth at the tooth surface and at the gums. This work contributes to our knowledge of bacterial diversity in the rat mouth.

## Introduction

Rats are the most frequently utilized rodent in periodontal disease pathogenesis research (nabr.org) and are used as a model for the human mouth. Despite the critical role of rat models in periodontal research, little information is available on the healthy oral bacteriome of the rat and its correlation with that of humans. Only two previous studies examined changes in the oral bacteriomes of rats due to dietary nitrate (Hyde et al., 2014), or ciprofloxacin dosing (Manrique et al., 2013) are available. Hyde and colleagues characterized the bacteriome of male Wistar rat tongues, to study the effect of dietary nitrate and compared their findings to humans (Hyde et al., 2014). They determined there were similarities in the bacteriome of the tongue between humans and rats, supporting the use of the Wistar rat model (Hyde et al., 2014). Manrique *et al*. studied the supragingival plaque of female Sprague-Dawley rats before and after ciprofloxacin dosing (Manrique et al., 2013). They found that supragingival plaque communities were dominated by *Streptococcus* and *Rothia* species (Manrique et al., 2013). Although these studies provided insight to a core bacteriome in the rat oral cavity, only a single location was examined for each study. The lack of understanding of the native oral microbiota in rats coupled with the significance of rat models in oral health and disease research highlights a critical gap in knowledge.

The goal of this study was to characterize the healthy rat oral bacteriome at the teeth and the toothless alveolar crest (i.e., gums) in 12 male Lewis rats. The first molar and the alveolar crest sample sites were chosen to further the understanding of the rat oral bacteriome at different tissue types, which are exposed to unique ecological factors (Mark Welch et al., 2020). We found that the two sample sites were not significantly different in diversity analyses, and that previously identified phyla in the rat oral cavity were also present in our study. Five taxa met our qualifications to be characterized as part of the core bacteriome, with the most frequent genera being *Streptococcus*, *Corynebacterium* 1, *Rothia*, *Haemophilus* and *Staphylococcus*. Of the 359 amplicon sequence variants (ASVs) detected in our study, six genera were enriched at the gums, and four genera and one order were enriched on the teeth. Although parallels between the human oral cavity and rats do exist, we found that the rat oral bacteriome is less diverse than the human oral bacteriome as reported in comparable human studies. We found that, although pooling swab samples from multiple rats generated more average reads per sample, analysis of individual swabs allowed for a more robust analysis of the bacteriome of an individual rat. This study establishes a baseline healthy bacteriome in the male Lewis rat for future comparison as it is the first study of its kind to analyze two different locations within the rat oral cavity. Furthermore, it demonstrates that this analysis can be done using single swabs from individual rats to obtain more robust information that may further demonstrate parallels between humans and rat oral cavities.

## Methods

### Animals for pooled swab analysis

Twelve male Lewis Rats (Charles River Laboratories, Wilmington, MA, USA), 10-12 weeks old, were housed in pairs and maintained at the University of Texas at Dallas Vivarium. Only males were used to eliminate variability that may be caused by female hormone cycles. The animals were housed in individually ventilated cages (IVC) with a 12-h light cycle and a 12-h dark cycle at 20-22.2°C. Rats ranged in weight from 250 to 325 grams and were fed dry Labdiet® Prolab RMH 1800 and sterile water *ad libitum*. The study was approved by the University of Texas at Dallas Institutional Animal Care and Use Committee (IACUC #16-05).

### Sample collection and DNA isolation from pooled samples

Rats were tranquilized with an intramuscular injection of 50-100 mg/kg ketamine hydrochloride and 20-50 mg/kg xylazine hydrochloride. Samples were collected from the first molar (tooth) and the alveolar crest (gum) on both sides of the rat mouth with Puritan HydraFlock swabs. Each side of the mouth was swabbed independently. Swabs were stored in dry storage tubes at −20°C. Samples collected on the same day from the same location (tooth or gum) and side (right or left) were pooled together into groups (3 rats per group; groups were named A, B, 1C, and 2C) to improve biomass recovery. This resulted in 16 pooled swab samples. Rat groups A and B, and groups 1C and 2C were collected on and extracted on the same days. DNA was isolated using the Qiagen DNeasy PowerSoil kit per the manufacturer’s instructions. Swabs were agitated for 8 seconds in the bead tube for biological sample removal. A total of 16 unique samples from 12 rats (8 from gums and 8 from teeth) were generated. The ZymoBIOMICS Microbial Community Standard, composed of multiple Gram-positive and Gram-negative bacterial species as well as two fungal species, was used as a positive control to determine the efficiency of the DNA isolation. The positive and negative controls were extracted with same DNA extraction kit; 75 μl volume of Community Standard or Qiagen DNA free water were utilized, respectively.

### Sample collection and DNA isolation from individual swab samples

Eight gum and tooth swabs from a single rat (H2) from a pilot study were extracted using the same methodology as pooled samples, however, only one swab was used for each extraction (n = 8). The male Lewis rat (Charles River Laboratories, Wilmington, MA, USA), was sampled twice between 10-12 weeks old, yielding 8 unique swabs.

### 16S rRNA V1-3 Sequencing

Samples were sequenced with Illumina MidSeq by Molecular Research Laboratory (Shallowater, TX). The V1-3 region of the 16S rRNA were amplified and sequenced. Primer sequences used were: bacteriome forward 27F (AGRGTTTGATCMTGGCTCAG) and reverse 519R (GTNTTACNGCGGCKGCTG). PCR conditions were as follows: initial denaturing at 94° C for 3 min, then 33 cycles of denaturing (94° C for 30 s), annealing (53° C for 40 s), elongation (at 72° C for 1 min), and a final elongation for 5 min at 72° C.

### Data analysis

All analyses were performed using QIIME2 (v. 2018.11 and v. 2020.2) (Bolyen et al., 2018). Raw Illumina sequences were imported as EMP Paired End Sequences and demultiplexed. Denoising was done with DADA2 (Callahan et al., 2016), which merged the reads, removed chimeras, and clustered features into amplicon sequence variants (ASVs). Truncation of forward and reverse reads was set such that the quality score for a nucleotide was no less than 18. The *phylogeny align-to-tree-mafft-fasttree* command was used to generate a rooted tree for diversity analysis. Diversity analysis was performed using the outputs from the *diversity core-metrics-phylogenetic* command. Alpha-diversity was calculated using the Shannon diversity index (H) and evenness score. Rarefication values were set to the lowest reads per sample; 7,065 was the value utilized for rarification in diversity analysis. The quantitative and qualitative unique fraction metric (unweighted and weighted UniFrac, respectively) was used to determine β-diversity. Bacterial taxonomy was assigned using the SILVA database (v. 132) (Glöckner et al., 2017; Pruesse et al., 2007; Quast et al., 2012; Yilmaz et al., 2014). *Cyanobacteria* was excluded as it is theorized to be a mitochondrial or chloroplast hit. All sequences characterized as unassigned or eukaryote were filtered out. Statistical testing was done using an ordinary one-way ANOVA for groups of four (n = 4), and two tailed paired t-test for groups of two (n = 8); significance was determined when *P* ≤ 0.05.

The R package decontam (Davis et al., 2018) was used to distinguish ASVs truly present in experimental samples from contaminating ASVs introduced from external sources. Given the relatively low biomass of the samples, the prevalence method was employed. This method calculates a chi-square statistic, expressed as a probability P, comparing the presence and absence of ASVs in true samples versus negative controls. Smaller values of P indicate a higher likelihood that the ASV is a contaminant. A threshold value of 0.5 was chosen; ASVs with values of P below this threshold were considered contaminants and removed from further analysis (Davis et al., 2018).

### Accession numbers

Sequence data generated in this study have been deposited into the Sequence Read Archive under accession numbers PRJNA875109.

## Results

### Bacterial amplicon sequence variant statistics for pooled rat groups

A total of 359 unique ASVs were identified at frequencies ranging from 1 to 113,816 across all 16 samples (Table 1). The sequence frequency per sample ranged from 7,065 to 39,208, and all samples were rarified to 7,065 sequences to improve even sampling for diversity analysis. The average length of the ASV sequences was 481.8 nt.

**Table 1.** Feature table for the bacteriome of pooled and individual swabs.

| *Microbiome* | *Reads per sample* |  |  | *ASV frequency* |  |  | *Total* |
| --- | --- | --- | --- | --- | --- | --- | --- |
|  | Min | Average | Max | Min | Average | Max | *ASVs* |
| **Pooled** | 7,065 | 20,820.7 | 39,208 | 1 | 927.9 | 113,816 | 359 |
| **Individual** | 6,665 | 15,026 | 20,916 | 3 | 703 | 33,813 | 171 |

### No significance difference in α-diversity among pooled rat groups or sample location

Comparison of rat groups without factoring in sampling location did not demonstrate a significant difference in number of ASVs detected, community evenness, or Shannon diversity (*P* > 0.05). Specifically, numbers of ASVs ranged from 22 to 104 (Fig. 1A), with an average of 49.6 (± 24.5) at the gums and 55.0 (± 12.2) at the teeth (Fig. 1B & Table 1). Evenness scores were not significantly different, ranging from 0.21 to 0.695 across all rat groups (Fig. 2A) and averaged 0.504 (± 0.158) at the gums and 0.499 (± 0.064) at the teeth (Fig. 2B). Shannon diversity scores (H) were not significantly different, ranging from 0.917 to 3.89 across all rat groups (Fig. 2C) and averaging 2.82 (± 0.983) and 2.85 (± 0.337) at the gums and teeth, respectively (Fig. 2D). 𝛽*-diversity analysis demonstrated increased diversity among gum compared to tooth samples* UniFrac measurements were utilized to determine β-diversity between samples. An unweighted UniFrac is a qualitative measure and only accounts for the presence or absence of an ASV, while a weighted UniFrac considers not only the presence or absence of an ASV but also its abundance. In the unweighted UniFrac, tooth samples clustered along the largest component of variance (25.25%, Axis 1), except for one of the 2C tooth samples (Fig. 3A). Although some gum samples appeared to cluster by rat group, there was no pattern of clustering across all gum samples (Fig. 3A). The weighted UniFrac axes were two times larger than the unweighted UniFrac axes, demonstrating increased diversity between samples when the abundance of ASVs was accounted for. Tooth samples clustered on the largest component of variance, apart from one group B sample (Fig. 3B). Gum samples overall did not cluster; however, Group A gum samples clustered tightly with one group B gum sample (Fig. 3B). This pattern is also observed between 1C and 2C gum samples. In summary, when only accounting for the presence or absence of an ASV, the samples were overall less diverse as demonstrated by the shorter axis. Larger diversity in the weighted UniFrac demonstrated that, although similar ASVs may be present among samples, the relative abundances increase the diversity between samples. Gum samples from group A and B also demonstrate tighter clustering, which could be due to external factors such as animals being housed together.

**Figure 1.**
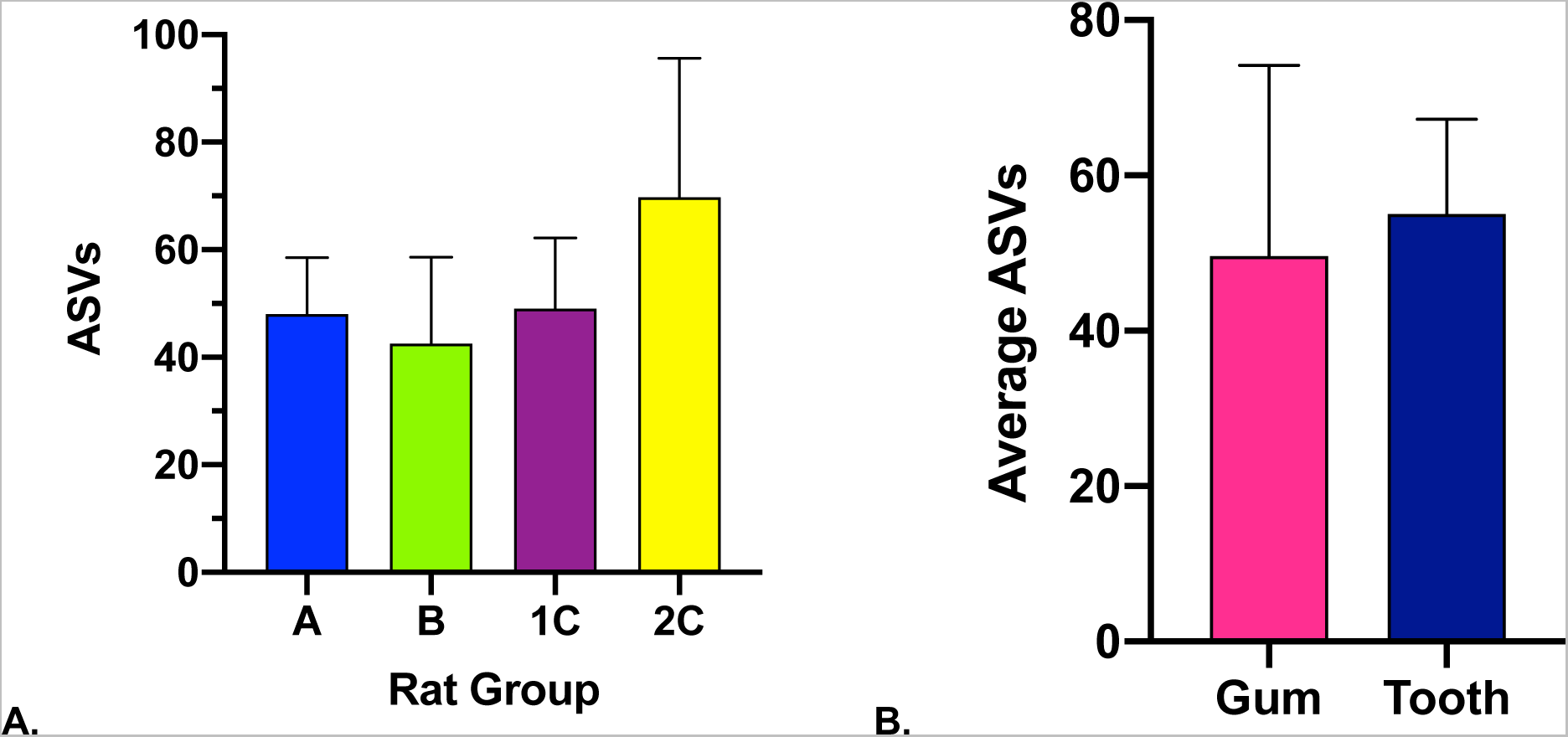
Observed bacterial ASVs in all rat groups (A) and collection locations (B). ASVs ranged from 22 to 104. There was no significant difference in the average ASVs for any rat group as determined with a one-way ANOVA. The average ASVs at the gums (49.63 ± 24.54) and teeth (55 ± 12.24) were not significantly different as determined with a two tailed paired t-test (n = 8).

**Figure 2.**
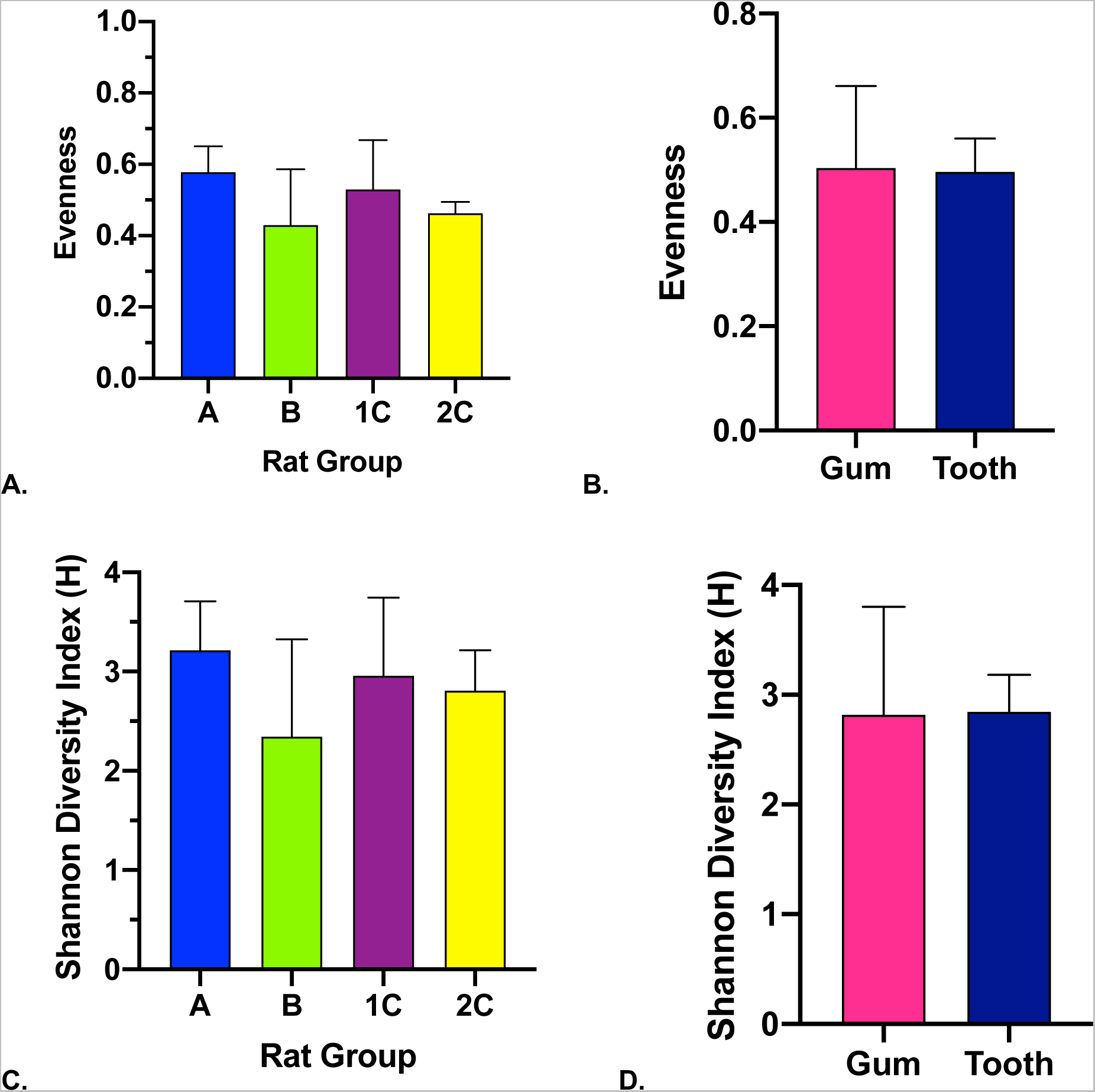
Alpha-diversity metrics across rat groups (A,C) and sample locations (B,D). Evenness scores (A-B) for bacteriome values closer to one indicate even community distributions and values closer to zero demonstrate uneven community distributions. No significant differences were found across any of the rat groups determined with an ordinary one-way ANOVA (n = 4). Average evenness at the gums (0.504 ± 0.158) and the teeth (0.499 ± 0.064) were not significantly different as determined with a two-tailed paired t-test (n = 8). Shannon diversity scores (H) (C-D) are shown for the bacteriome across rat groups and sample locations. There were no significant differences between rat groups (n = 4) determined with a one-way ANOVA. A two-tailed paired t-test did not find a significant difference between the averages at the gums (2.82 ± 0.983) or teeth (2.85 ± 0.337).

**Figure 3.**
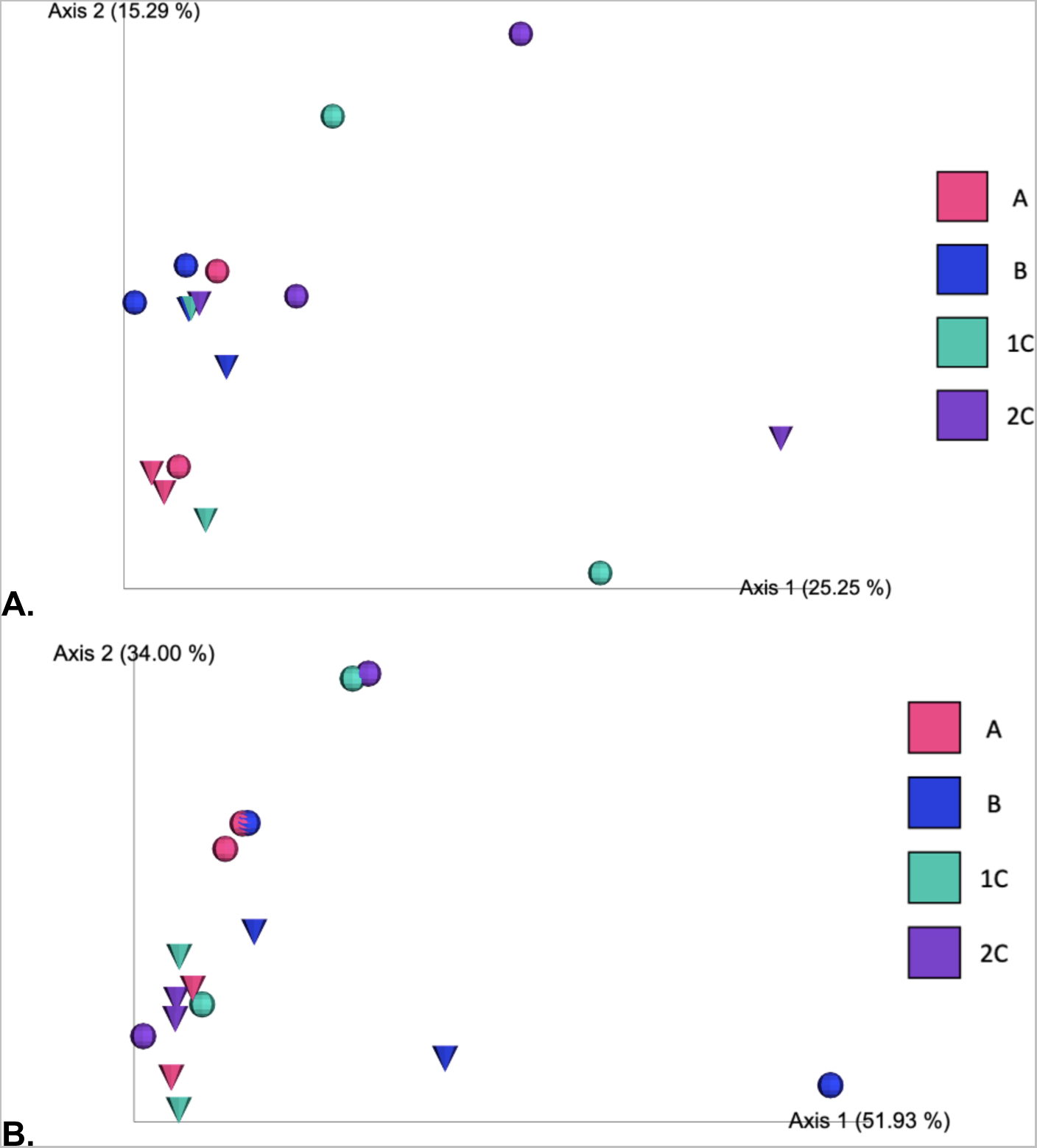
Bacteriome beta-diversity metrics using Unweighted (A) and Weighted (B) UniFrac. Gums are shown as spheres and teeth are demonstrated as cones. When accounting for the presence or absence of an ASV (A), tighter clustering can be observed for some of the tooth swabs indicating there are similar ASVs present at this location across rat groups. The presence of gum samples near the tooth cluster indicates it is unlikely there is a clear difference in the species present at teeth and gums. When ASV presence and abundance are considered (B), tooth samples cluster on the largest component of variance, with the exception of one of the group B tooth samples. All tooth samples cluster on the second largest component of variance. Half of the gum samples cluster with one or two other samples from their rat group but there is not tight clustering based on sample location for gum swabs. This indicates that the relative abundance of the present ASVs is similar across the clustering samples.

### Taxonomy at the Phylum level supports previous reports

Taxonomy results to the Phylum level were assessed to better compare to previously published reports (Hyde et al., 2014; Manrique et al., 2013). At the Phylum level, the majority of hits were associated with *Firmicutes*, *Actinobacteria* and *Proteobacteria*. *Bacteroidetes* were identified in 11 of the 16 samples, ranging from 3 to 1,270 reads (Supplemental Fig. 1). One of the group B gum samples showed an increased prevalence of *Proteobacteria*, which was correlated to enrichment in the genus *Methylobacterium* in downstream analysis (Fig. 4 B & C).

**Figure 4.**
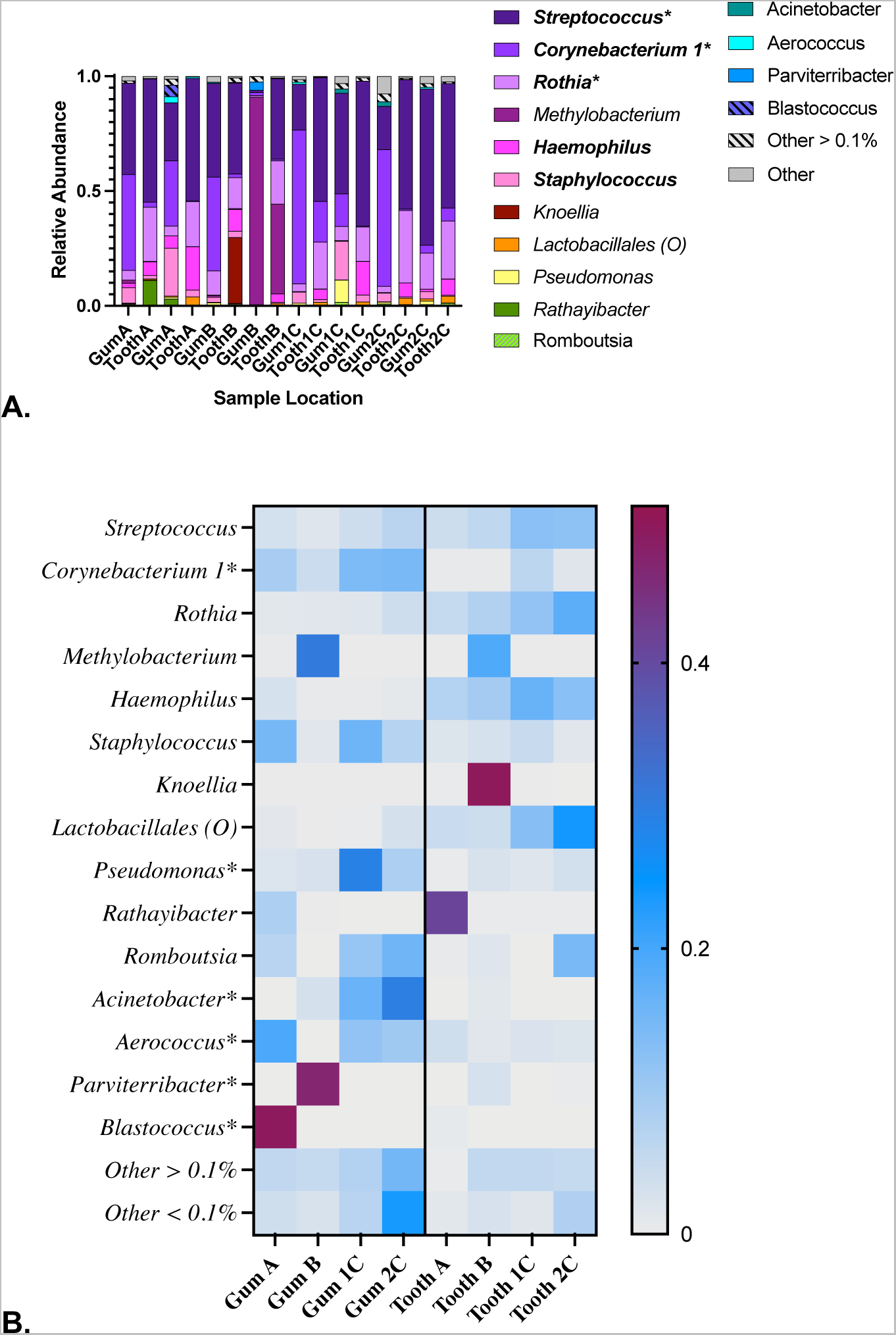
Core bacterial enrichment and taxonomy, including taxa contributing 1% or greater to the total reads. (A) Taxonomy bar plot showing the most abundant taxa present across all samples after removal of contaminants with DECONTAM with an A value of 0.5. The relative abundances are calculated by determining the percent of each taxonomy within a sample, generating bar plots that total to one. Bolded genera were detected in all samples and genera with an * contribute 10% or greater of the total reads (B) Heat map of taxa contributing 1% or greater to the total reads demonstrating enrichment at the teeth or gums. Samples from the gum or teeth were combined for each rat group and the averages of those are presented here. Each taxon shown except *Lactobacillales* (Order) corresponds to a genus. Taxa with an * were enriched at the gums. The genera *Rothia*, *Haemophilus*, *Knoellia*, *Rathayibacter* and the order *Lactobacillales* were enriched at the teeth.

### Five taxa met the core bacteriome criteria defined in this study

In this study, the core rat bacteriome was defined as taxa that accounted for ≥1% of the total reads across all samples. Of the 359 unique ASVs, only five taxa met the criteria to be characterized as part of the core bacteriome. The genus *Streptococcus* was ubiquitous (i.e. present in every sample), accounting for 42.3% of the total reads across all samples (Fig. 4A). Additionally, the genera *Corynebacterium* 1, *Rothia*, *Haemophilus* and *Staphylococcus* were ubiquitous, respectively contributing 17%, 14.7%, 4.6%, and 3.5% to the total reads. Of the genera present in all samples, *Streptococcus*, *Corynebacterium* 1, and *Rothia* accounted for 10% or more of the total reads across all samples.

We determined whether specific taxa were enriched at the tooth or gums. Enrichment was reached when at least 80% of the total reads for a specific taxon were associated with either the teeth or gums. Of the taxa contributing to ≥1% of the total read frequency, six genera were enriched at the gums (*Corynebacterium* 1, *Pseudomonas, Acinetobacter, Aerococcus, Parviterribacter* and *Blastococcus*), and four genera and one order (*Rothia*, *Haemophilus*, *Knoellia, Rathayibacter* and the order *Lactobacillales*) were enriched on the teeth (Fig. 4B). Of the core oral bacterial taxa, neither *Streptococcus* or *Staphylococcus* demonstrated enrichment at the gums or teeth. Although contributing <1% to the total reads, the genera *Jeotogalicoccus*, *Modestobacter*, and *Bacteroides* were found at ≥80% frequency on the gums; *Xanthomonas* and *Cloacibacterium* were found exclusively on the gums; and the genera *Williamsia, Klebsiella* and *Granulicatella* were found exclusively on teeth samples.

For some taxa, one rat group accounted for 93-100% of the total reads. This was observed most frequently in three of the group B samples. One rat group B gum sample corresponded to ≥60% of the total reads for *Methylobacterium*, *Parviterribacter*, and *Modestobacter*. The tooth sample from the same side of the mouth accounted for 37% of the *Methylobacterium* reads. In the same rat group (B), the tooth sample from the opposite side corresponded to 99% of the *Knoellia* and all the *Williamsia* reads. This may explain the observed clustering in the unweighted UniFrac and the loss of clustering in the Weighted UniFrac for this rat group (Fig. 3AB).

### Individual swab analysis

Eight swabs obtained from a single rat from an initial pilot study were chosen to compare if the same analysis would be possible using individual swabs instead of pooling swab samples. Table 1 shows the comparison of features and frequency data for pooled (n = 16) and individual (n = 8) swabs. A total of 207 unique ASVs were identified with a total frequency of 120,208 across all samples (Table 1). The reads per sample ranged from 6,665 to 20,916 with an average frequency of 15,026 (Table 1). The average nucleotide length of the paired joined reads was 483.24 nt.

### Pooled and single swab samples have overlapping taxa present

Taxonomy analysis between the pooled samples and the individual swabs showed parallels between the data collection methods. Similar genera were identified in both data sets including, *Streptococcus*, *Rothia*, *Lactobacillales* (O), and *Haemophilus* (Fig 4A & 5A). Of the 171 unique ASVs across the single swab samples, 10 taxa were present at ≥1% across all samples. The genera *Streptococcus*, *Ralstonia* and *Cutibacterium* were ubiquitous, accounting for 54.2%, 8.8% and 3.4%, respectively, of the total reads across all samples (Fig. 5A). Enrichment was reached when at least 80% of the total reads for a specific taxon were associated with either the teeth or gums. Of the taxa contributing to ≥1% of the total read frequency, three genera (*Alishewanella*, *Cutibacterium*, and *Veillonelia*) were enriched at the gums, and one family (*Aerococcaceae*) was enriched on the teeth (Fig. 5B). We conclude that the oral bacteriome can be elucidated using analysis of single swabs.

**Figure 5.**
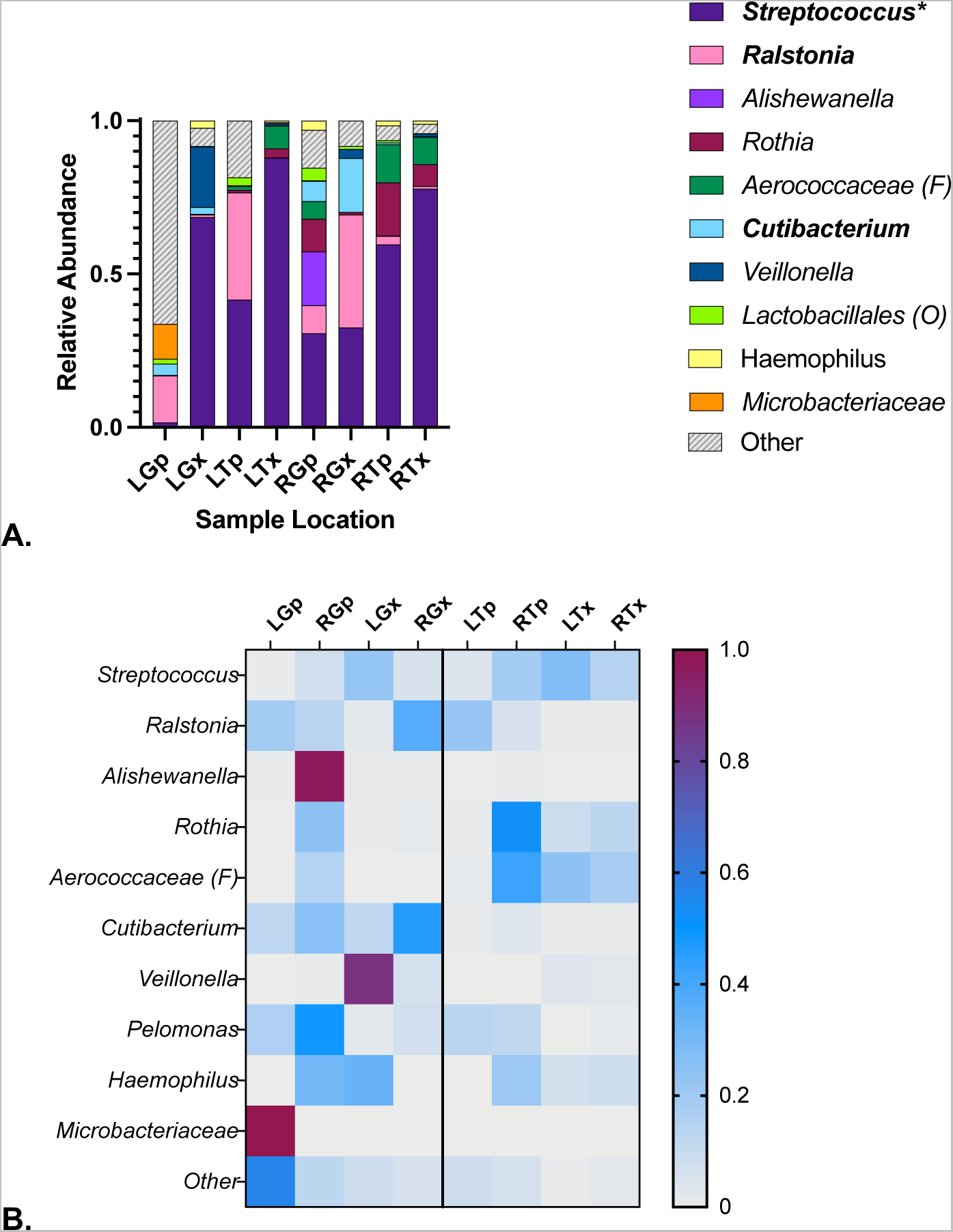
Single swab taxa contributing 1% or greater to the total reads across all samples. (A) Bar plot for taxa contributing 1% or greater to the total reads across all samples, bolded taxa are present across all samples, * indicate the taxa contributed 10% or greater to the total reads across all samples. As observed in pooled samples *Streptococcus*, *Rothia* and *Haemophilus* are present in single swab samples. (B) Heat map demonstrating enrichment between taxa either the teeth or gums. Three genera, *Alishewanella*, *Cutibacterium*, and *Veillonelia,* were enriched at the gums, and one family: *Aerococcaceae* was enriched on the teeth.

## Discussion

Rats are the most common animal model to study periodontitis disease and progression (Struillou et al., 2010), yet only two studies are currently available that examine the oral bacteriome of the rat (Hyde et al., 2014; Manrique et al., 2013). Periodontitis is a disease that results in the loss of a tooth and is caused by bacterial contamination (Marcenes et al., 2013). Severe periodontitis is estimated to affect 11% of the world population (Marcenes et al., 2013). The lack of information available motivated the exploration and characterization of core members of the bacteriome at two unique oral niches in the healthy male Lewis rat. This study represents the first of its kind to examine two unique locations within the rat oral cavity with the aim of defining the core oral bacteriome using 16S rRNA sequence analysis. This information sets a baseline for the healthy rat oral cavity and will be utilized as a guide for future studies seeking to identify changes in the rat oral bacteriome linked to biomaterials implantation, use of novel dental implant materials, surface treatments, and dental implant coatings (Gindri et al., 2014, 2016).

### Amplicon sequence variants

In this study we utilized ASVs due to the improved resolution over the previously utilized operational taxonomic units (OTUs) (Callahan et al., 2017). Utilizing ASVs instead of OTUs allows for comparisons between samples that utilize the same primer sets but were not sequenced and analyzed together. In this study, we identified 50.0 (± 12.2) ASVs at the tooth and 49.6 (± 24.5) ASVs at the gums (Table 1). A previous report characterizing the rat tongue flora in healthy male Wistar rats identified an average operational taxonomic unit (OTU) of 99.3 ± 40 (Hyde et al., 2014). Based on topography and tissue type, we compared our gum samples to this study and found that the rat gums are less diverse than the tongue. A study examining the supragingival plaque in 6-month-old female Sprague-Dawley rats identified 575 OTUs (Manrique et al., 2013) indicating that the dental plaque is occupied by more phylotypes than the tooth in this study. However, the differences between rat strain, diet, housing, method of collection and utilizing OTUs instead of ASVs may contribute to the variability noted here.

### Diversity at the alveolar crest and first molar were not significantly different

We examined diversity at the tooth and gum in the healthy male Lewis rat oral bacteriome to determine if sample location influenced diversity. Alpha-diversity metrics to examine the diversity within the sample were analyzed with both Shannon diversity and evenness metrics. We did not observe significant differences in alpha-diversity between the two sample locations or rat groups (Fig. 2). Hyde (Hyde et al., 2014) and colleagues found similar Shannon diversity values at the tongue (H = 2.92 ± 0.53) compared to the averages reported here at the gums (2.82 ± 0.983) or teeth (2.85 ± 0.337) (Fig. 2 C-D). This may demonstrate that these three niches exhibit similar ecological richness, and although the tongue exhibited higher OTU values, Shannon diversity accounts for both evenness and abundance. With respect to this metric, the larger amount of OTUs is not directly correlated to higher alpha-diversity values.

### Samples do not cluster by collection site

Examining between-sample diversity was done using two phylogenetic UniFrac measurements, unweighted (qualitative), and weighted (quantitative) (Fig. 3 A-B). The unweighted UniFrac measured diversity by the presence or absence of ASVs. Unexpectedly, the samples did not cluster by sample location (Fig. 3A). Teeth samples did demonstrate improved clustering compared to gum samples across both axes, however, on the largest component of variance (Axis 1 25.25%) discriminated one of the Group 2C tooth samples from all other tooth samples. This demonstrates that many of the tooth samples share similar ASVs, yet one of the Group 2C tooth samples does not share many of these ASVs. Gum samples for groups A and B appear to cluster with samples from their own group across the second greatest component of variance (axis 2 15.29%). ASVs present on the teeth and gums in these groups are similar across all sample locations.

Accounting for the abundance of ASVs in the weighted UniFrac resulted in marked differences from the unweighted UniFrac (Fig. 3B). The components of variance were markedly increased in the weighted UniFrac demonstrating larger diversity between samples. Overall, tooth samples had improved clustering along both axes apart from group B tooth samples. Group B showed the largest variance, and this may be explained by the presence of *Methylobacterium* present at 60.8% and 37.3% in a gum B and tooth B samples, respectively (Fig. 3B).

### Taxonomy analysis

From this study, we discovered 15 genera that constitute >1% of the total detectable taxa. Our study focused on the microbiota at two locations, the tooth surface and toothless alveolar crest. We identified the presence of the phyla *Firmicutes*, *Actinobacteria, Proteobacteria* and *Bacteroidetes* (Fig. 4A). Additionally, the genera *Streptococcus*, *Corynebacterium*, *Haemophilus*, and *Staphylococcus* were identified at the tooth and gum as shown in Figure 4. These findings are supported by the previous report of the rat tongue and supragingival plaque microbiota (Hyde et al., 2014; Manrique et al., 2013). However, our study identified the presence of, *Methylobacterium*, *Knoellia*, and *Pseudomonas* as shown in Figures 4 A and B, which were not previously reported (Hyde et al., 2014; Manrique et al., 2013). The findings illustrated in Figure 4 A and B support the characterization of rat plaque bacteriome in which *Streptococcus* and *Rothia* were major contributors to the present taxa (Manrique et al., 2013). Together these findings demonstrate the differences between the taxa found among different rat oral niches, indicating that, similar to humans, different phylotypes preferentially colonize specific niches.

### Comparison between the human oral bacteriome and Lewis rats

After identifying a core bacteriome of the healthy male Lewis rat, we identified the similarities with the healthy human oral microbiome. Studies have reported over 700 different species of bacteria within the nine oral niches in humans; some species preferentially colonize certain niches that vary in topography and chemistry (Aas et al., 2005; Dewhirst et al., 2010; How et al., 2016; Mark Welch et al., 2020; Zhang et al., 2018). In this analysis, we looked at gum and tooth sites because a previous study demonstrated that some genera like *Streptococcus* are found across all oral niches, while others such as *Actinomyces* preferentially colonize surfaces such as teeth in the human oral cavity (Aas et al., 2005). Although humans do not have a toothless alveolar crest (as rats do), we compared our findings to the bacteriome of soft tissue in humans, which include the gums, cheeks and hard palate, which have been found to have similar microbial compositions (Mark Welch et al., 2020). There was not a significant difference between Shannon diversity scores for the tooth or gum sample sites (Fig. 2D). Shannon diversity scores at the rat tooth and gums (2.85 ± 0.337 and 2.82 ± 0.983 respectively) are lower than at the human tongue (H = 5.536 ± 0.362) (Hyde et al., 2014). This is consistent with a previous study reporting lower diversity on the rat tongue with a reported Shannon diversity value of 2.92 (± 0.53) (Hyde et al., 2014). Examination of the supragingival plaque biofilms in the oral cavities of healthy children identified a Shannon diversity score of 3.793 (mode) which was significantly greater compared to a dental caries (diseased) subset (Espinoza et al., 2018). However, comparison between peri-implantitis and healthy subgingival plaques indicated higher alpha-diversity in peri-implantitis with a greater amount of Gram-negative species (Koyanagi et al., 2010). Overall, the alpha-diversity of the rat oral bacteriome appears to be less complex than human oral cavities (healthy or diseased). This may be correlated to the differences in diet and oral hygiene habits between humans and rats.

In this study, we identified four phyla in the healthy rat oral cavity: *Firmicutes* (genera *Streptococcus, Staphylococcus* and the order *Lactobacillales*), *Actinobacteria* (genera *Corynebacterium 1, Rothia,* and *Knoellia*), *Proteobacteria* (genera *Methylobacterium, Haemophilus* and *Pseudomonas*), and *Bacteroidetes*. Comparatively, in humans, there are six different phyla that dominate the human oral bacteriome including *Firmicutes*, *Actinobacteria*, *Proteobacteria*, *Fusobacteria Bacteroidetes* and *Spirochaetes* identified through 16S rDNA profiling (Dewhirst et al., 2010; Verma et al., 2018). The rat subjects examined here were dominated by *Firmicutes* (*Streptococcus*, and *Staphylococcus*) (Fig 4A). We found that *Streptococcus* was the most abundant genus in the rat oral cavity which is also the dominant genus in the healthy human oral cavity (Dewhirst et al., 2010). We identified that *Haemophilus* and *Streptococcus* contributed greater than 1% of the total reads and were found across all samples (Fig. 4 A). Hyde and colleagues compared human and rat tongues and found *Haemophilus* and *Streptococcus* to be among the most common genera in humans and rats (Hyde et al., 2014). From these findings there are parallels between rats and human oral cavities, however, further studies need to be carried out to further elucidate direct parallels.

### Limitations of study

Our study has limitations. (1) Taxonomic classification was only to the genus level. Further elucidation to the species level would improve the characterization of the rat oral bacteriome and allow for improved comparison to the human oral cavity. (2) Pooling samples was initially done to ensure enough DNA was present to carry out the analysis, however, the limitation of this methodology was demonstrated with group B rat samples, which were outliers due to *Methylobacterium* abundance. This result led us to attempt analysis with single swabs from an individual rat. Although fewer total ASVs were identified (Table 1), we identified similar taxa among the individual rat mouth swab samples (Fig. 5A) and the pooled swabs (Fig. 4A). This suggests that samples do not require pooling to achieve sufficient biomass. We note that use and analysis of appropriate negative controls are critical for low biomass samples; in this study we used decontam for this purpose (Davis et al., 2018). (3) Rarification values are only utilized for diversity analysis, so although individual swabs have lower average reads per sample, this does not interfere with the taxonomy classification. However, due to rarification, a large portion of sequences are not included in the diversity analysis. This may have influenced clustering in the UniFrac analysis. (5) Characterizing the rat oral mycobiome would improve understanding of the rat oral cavity (Falsetta et al., 2014; Peters et al., 2017; Zaura et al., 2009). (6) Both female and male rats should be used to determine whether differences exist in their oral microbiota.

## Acknowledgments

This work was supported by National Institutes of Health grant R01DE026736 to D.R. and the Cecil H. and Ida Green Chair in Systems Biology Science to K.P.

## Supplemental Figures

**Figure S1.**
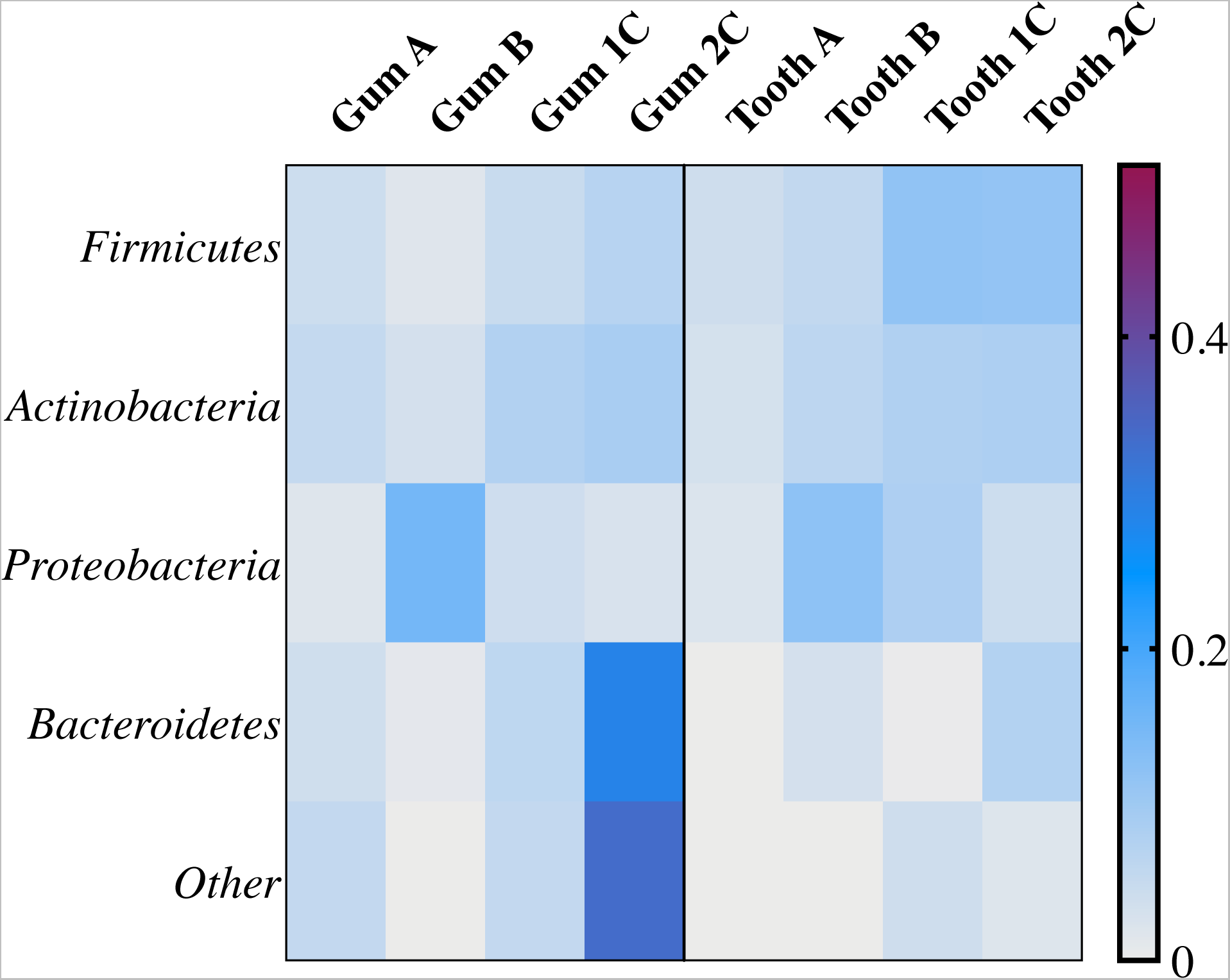
The phyla of bacteria identified across all rat samples and locations. The heat map represents the spread of each Phylum across all samples in which each row totals to one. The “Other” row represents taxa that contribute less than 0.001% of the total features identified across all samples. Gum 2C demonstrates the largest amount of taxa from the other category corresponding to the genus Chloroflexi which is found exclusively on one of the 2C gum samples.

